# Semi-supervised machine learning facilitates cell colocalization and tracking in intravital microscopy

**DOI:** 10.1101/829838

**Authors:** Diego Ulisse Pizzagalli, Marcus Thelen, Santiago Fernandez Gonzalez, Rolf Krause

**Affiliations:** Faculty of Biomedical Science, USI, Institute for Research in Biomedicine, CH6500 Bellinzona – Switzerland; Institute of Computational Science, USI, CH6900 Lugano – Switzerland

**Author notes:** Co-last and corresponding authors and.

**Keywords:** Colocalization, Segmentation, Tracking, intravital microscopy, Semi-supervised learning

## Abstract

2-photon intravital microscopy (2P-IVM) is a key technique to investigate cell migration and cell-to-cell interactions in organs and tissues of living organisms. Focusing on immunology, 2P-IVM allowed recording videos of leukocytes during the immune response, highlighting unprecedented mechanisms of the immune system. However, the automatic analysis of the acquired videos remains challenging and poorly reproducible. In fact, both manual curation of results and tuning of bioimaging software parameters among different experiments, are required. One of the most difficult tasks for a user is transferring to a computer the knowledge on what a cell is and how it should appear with respect to the background, other objects, or other cell types. This is possibly due to the low specificity of acquisition channels which may include multiple cell populations and the presence of similar objects in the background.

In this work, we propose a method based on semi-supervised machine learning to facilitate colocalization. In line with recently proposed approaches for pixel classification, the method requires the user to draw some lines on the cells of interest and some line on the other objects/background. These lines embed knowledge, not only on which pixel belongs to a class or which pixel belongs to another class but also on how pixels in the same object are connected. Hence, the proposed method exploits the information from the lines to create an additional imaging channel that is specific for the cells fo interest. The usage of this method increased tracking accuracy on a dataset of challenging 2P-IVM videos of leukocytes. Additionally, it allowed processing multiple samples of the same experiment keeping the same mathematical model.

## Introduction

2-photon intravital microscopy (2P-IVM) is a key technique to investigate cell-to-cell interactions in organs and tissues of living animals [1]. Focusing on immunology, the analysis of 2P-IVM videos revealed unprecedented mechanisms of the immune system such as novel interaction patterns between immune cells, host, and pathogens [2].

The standard imaging pipeline involves the acquisition of 4D videos (3D volumes at different time points) in a properly prepared sample. Once videos have been acquired, they are analyzed by performing cell detection, cell tracking, and computation of track-based measures, which are used to quantify cell migration and interaction (Figure 1). 2P-IVM requires a fluorescent sample. Although certain molecules of living organisms are capable to spontaneously emit fluorescence, like, for instance, the collagen fibers, the great majority of the cells and tissues need to be labelled with fluorescent markers to be visible in 2P-IVM acquisitions. The most common labelling methods are genetically modified (GM) animals, labelling *in vitro* with fluorescent dies, usage of fluorescent antibodies. When multiple cell populations are imaged, to distinguish between them it is desirable to acquire multiple acquisition channels having each channel specific for one cell type. This involves the usage of different fluorescent labeling (each with a specific emission wavelength), in combination with a set of optical filters. Such a combination of labeling and filter set is critical to distinguish a cell population from another, and from the background.

**Figure 1.**
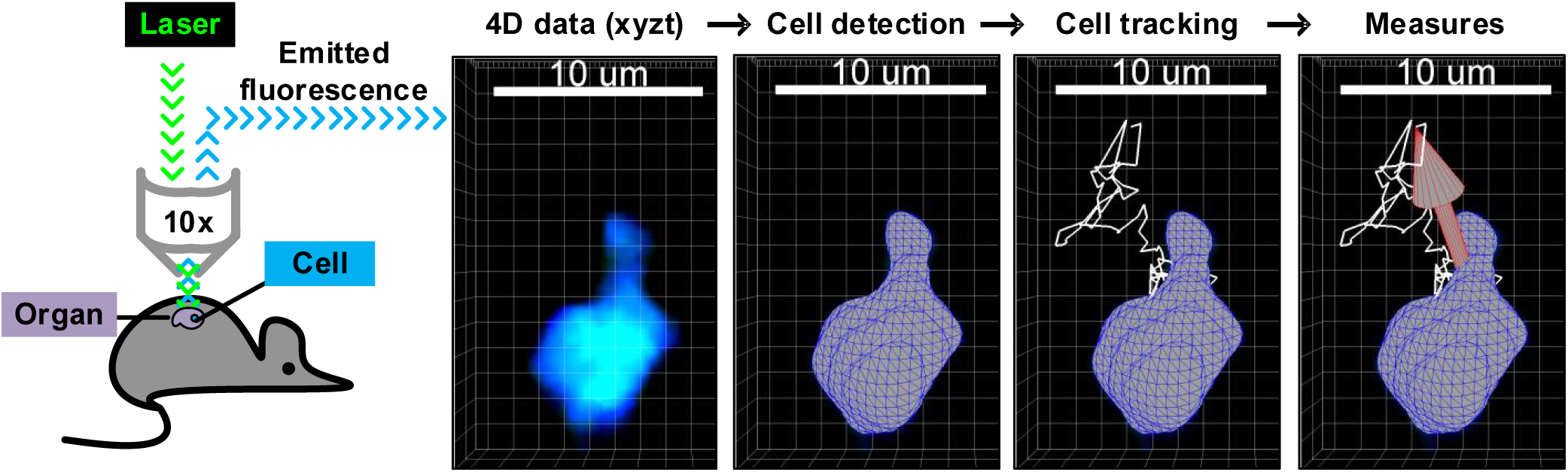
2P-IVM imaging pipeline. A laser stimulates fluorescent cells within an organ. Then, emitted fluorescence is converted into 4D videos (3d stacks at different time points). Cells are detected by separating fluorescent intensity above a threshold from background. Cell tracking happens by linking cells at different time points. Finally, measures on cell tracks are computed.

Indeed, if cells are poorly visible, or similar to other objects, errors in cell detection can be introduced. These can subsequently compromise tracking and affect the final measurements.

Unfortunately, fluorescently labeled immune cells often appear in more than one acquisition channel or exhibit bizarre appearance. This is due either to the broad emission spectrum of the used fluorescent markers (Figure 2, A) or to the incorporation of fluorescent material acquired, for instance, via phagocytosis. The effect of phagocytosis is often visible when phagocytic cells (i.e. neutrophils) are labeled *in vitro* with the formation of bright spots in the lysosomes due to the accumulation of the marker (Figure 2, B).

**Figure 2.**
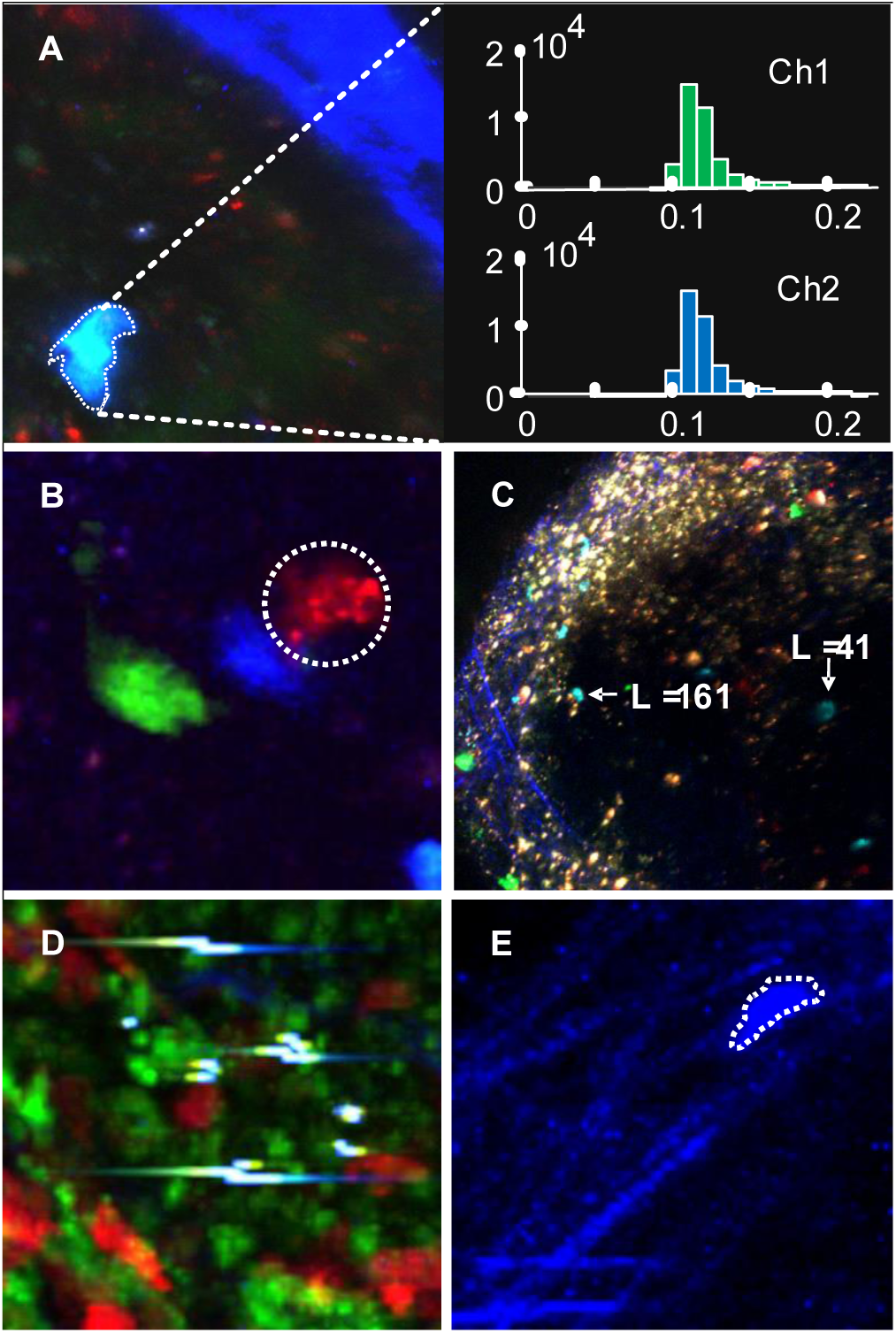
Challenges in 2P-IVM imaging of immune cells. **A**. CFP neutrophils emitting fluorescence in two acquisition channels with similar intensities. **B.** Neutrophils constitutively expressing GFP (green), CFP (blue) or labelled in vitro with CMTMR die (red). The CMTMR labelled cell shows non-uniform brightness and accumulation of fluorescent die in the lysosomes. **C.** Non spatially-uniform brightness showing a decrease of luminance (L) towards the bottom-right part of the image. **D.** Imaging artifacts due to saturation which were introduced by the movement of the animals and surgery defects. **E.** CFP neutrophil with elongated shape migrating on collagen fibers.

Additionally, the fluorescence emitted by cells undergoes diffraction throughout the sample, introducing a brightness variation in space (Figure 2, C). Brightness variation and artifacts can further be introduced by photo-damage and insufficient isolation of animal movements (Figure 2, D). Lastly, auto-fluorescent objects, such as background and collagen fibers can appear in the same acquisition channels of cells with similar intensity (Figure 2, E).

Therefore, specific image processing techniques are required to generate specific channels where only the cells of interests are visible.

However, the multiple aforementioned challenges hamper the usage of generic colocalization methods. Brightness thresholds to separate background from the cells of interest should be adapted among different areas of the same acquisition and different experiments. This reduces the usability of specialized imaging software packages such as Imaris (Bitplane inc.) and introduces research bias.

Although morphological information can in principle be taken into account to distinguish immune cells from the background, their high plasticity and commonly non-convex shapes introduce additional challenges to consider all the possible cases.

In this work, we propose an imaging processing method based on semi-supervised machine learning. This approach facilitates the transmission of knowledge on the appearance of cells from an imaging expert to a computer, in line with recently proposed methods for clustering [3]. This knowledge is then exploited to create an additional imaging channel which is more specific for the cells of interest. The usage of this channel improved tracking accuracy on challenging 2P-IVM videos from the Leukocyte Tracking Database [4]. Additionally, the proposed method allows using the same mathematical model to analyze different samples of the same experiment.

These are for instance videos from samples labeled using the same markers, but at different time points or in different animals. By providing examples of desired cells across all the samples, the computer automatically identifies the common parameters for the analysis. Hence, it does not require to adapt brightness thresholds from experiment to experiment, with a positive impact on the reproducibility of results.

The application of the method is supported by a user-friendly software with a graphical interface that allows selecting which features to use according to the videos to be analyzed. Such software is available as a plugin for Imaris at http://www.ltdb.info/tools.

## Results

### Color features improve tracking of poorly visible cells

The method was applied on the video 10 from LTDB, which included B cells labelled with CTV (red) migrating in the spleen of a mouse with a green-autofluorescent background (Figure 3, A). Although the signal to noise ratio of the video is very high (¿ 43) and the majority of the cells appear bright, their brightness can vary over time because of the challenges previously described. This variability introduces tracking errors when cells temporarily appeared with a brightness level similar to the background, interrupting tracks and creating multiple track fragments of short duration (Figure 3, B, C blue). By contrast, the proposed method learned to distinguish poorly visible cells from the background. By using the imaging channel generated by the method, tracks were computed without errors (Figure 3, B, C yellow).

### Color features improve separation of CFP cells from SHG background and GFP

Certain fluorescent cells are visible in multiple acquisition channels. The proposed method was applied to analyze the movement of neutrophils, adoptively transferred from a CK6/ECFP animal [5] to a CD11c/GFP animal [6] (Figure 4, A). Additionally, the acquired data included viral particles labeled with the DiD fluorescent die [7].

**Figure 3.**
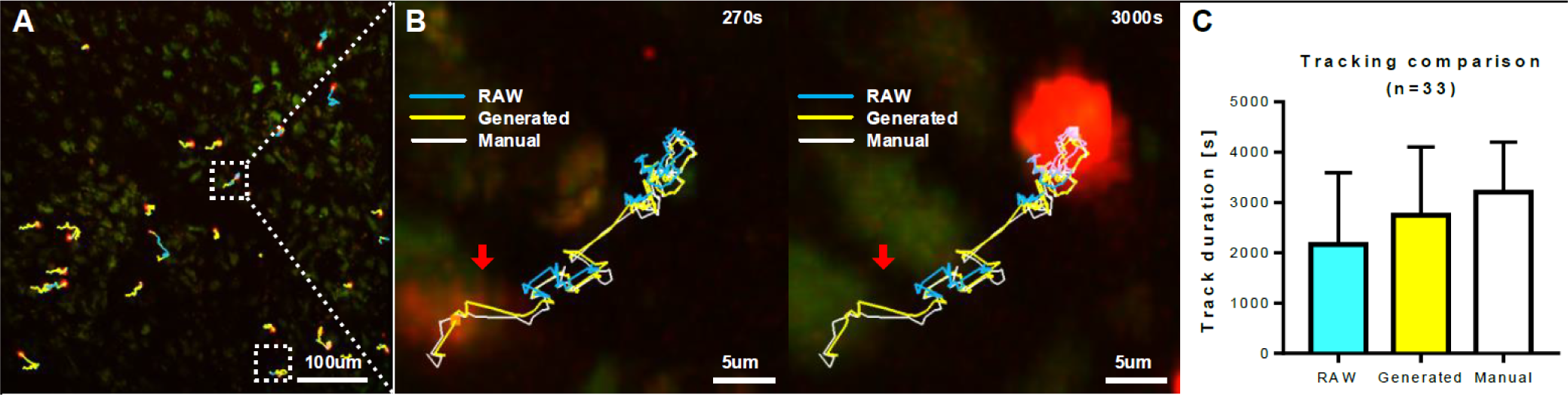
Tracking improvements for poorly visible cells. **A.** 2P-IVM micrograph showing CTV B cells (high red) in the spleen, migrating over an autofluorescent background visible (green / low red). **B** Sequence of micropgraph at two time points, showing a cell appearing with different brightness levels. Lines are the tracks obtained with different methods, showing errors of automatic tracking using RAW data (blue line) when a cell appeared with low brightness (red arrow). By contrast, the tracks obtained using the generated colocalization channel (yellow line), correspond to manual tracking (white line). **C.** Comparison of the track duration on the entire video.

**Figure 4.**
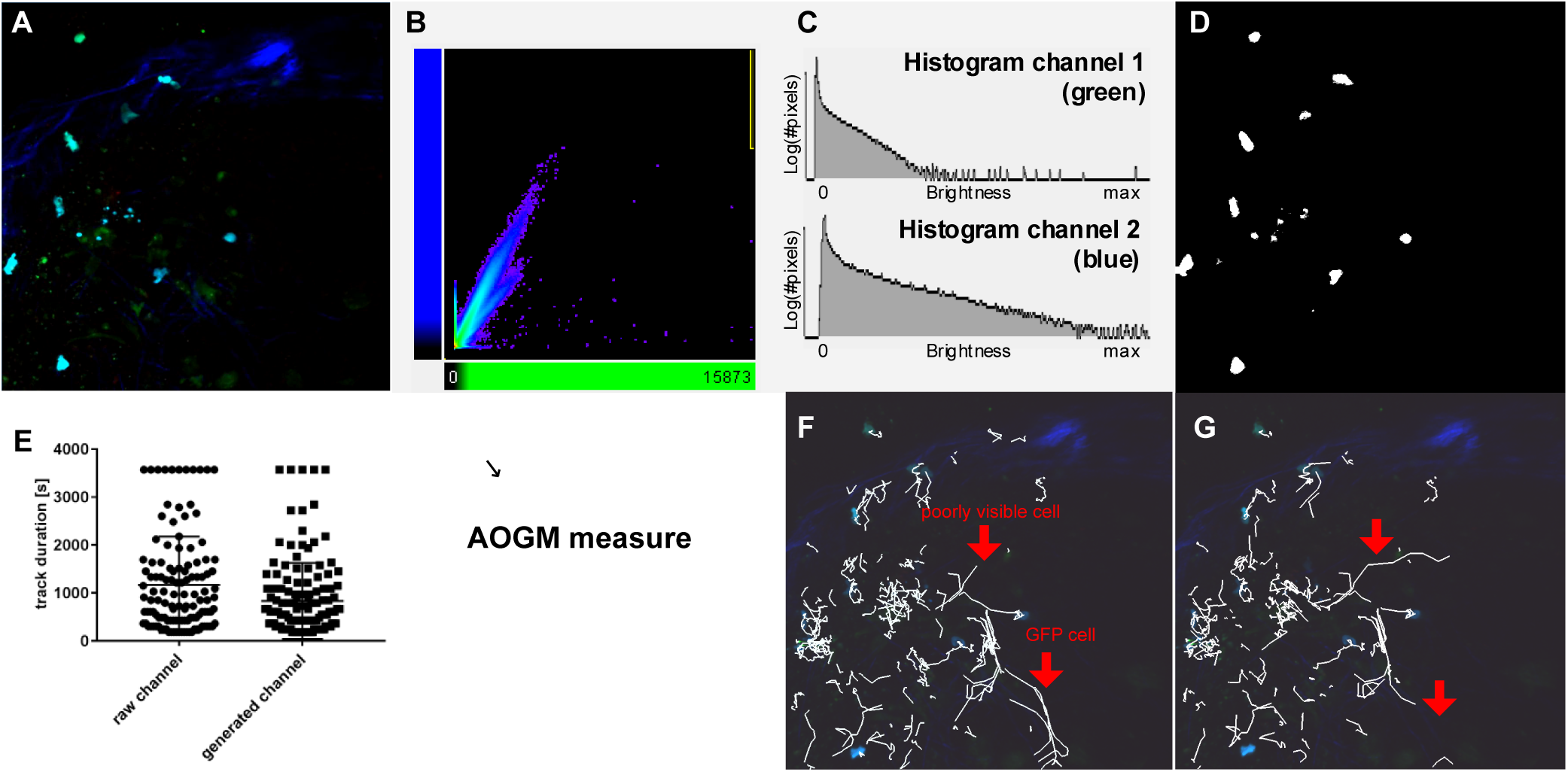
Pixel-wise classification. **A.** 2P-IVM micrograph showing CFP neutrophils (light blue) in the popliteal lymph node of a CD11c/GFP animal (green) with collagen structures (blue – SHG) and viral particles (red – DiD). **B** Scatter plot showing the intensity of each pixel in two acquisition channels. The central tails are cells appearing in more than one imaging channel. **C.** Histogram of pixel intensity in two acquisition channels. **D.** Results (binary mask) of pixel classification. **E.** Comparison of track duration when using the raw channels or the generated channel. **F.** Tracks (white lines) obtained using the original imaging channels including errors (red arrows). **G.** Tracks (white lines) obtained using the generated colocalization channel specific for CFP cells.

In the videos acquired via 2P-IVM, the emitted spectrum of neutrophils was captured by two imaging channels centered on the green and blue wavelengths. However, these channels captured also the fluorescence from the CD11c/GFP cells and the auto-fluorescence of collagen (second harmonic generation). The result was an overlap of the fluorescence emitted by the two cell populations and collagen (Figure 4, B) which resulted in poorly separable distributions (Figure 4, C). Additionally, the acquired videos exhibited spatially non-uniform brightness and brightness variation over time.

To separate neutrophils from the other cells and collagen fibers, each pixel was described using a set of features including the color of the pixel, the average color in a small circular neighborhood of radius = 3*µm*, and the average color in a larger circular neighborhood of radius = 7*µm*.

20 annotations were added by an imaging expert on the cells of interest, and 20 annotations on the other cells/background. These annotations were used to classify every single pixel, creating an additional imaging channel which is specific for the neutrophils (Figure 4, D).

The usage of the computed colocalization channel improved the accuracy of cell tracking (Figure 4, E) excluding unwanted objects such as not-motile pieces of collagen fibers and GFP cells (Figure 4, F), and improving the detection of poorly visible cells (Figure 4, G).

Note for the preprint version: In this version, we quantify only track duration using the raw channel and using the colocalization channel. This is a weak indicator of tracking accuracy. We will provide improved quantification, using the AOGM measure with respect to the ground truth in the next revised version. Additionally, we will include screenshots of the tracks.

### Path-features separate cells from cell debris

The annotations provided by the user in form of a line (Figure 5, A, yellow and red lines) were exploited, not only to detect pixels belonging to foreground or background, but also to decompose images into superpixels (Figure 5, A, white lines). Considering a cell as a group of pixels, a line drawn between the pixels of the cell contains information on how these pixels are connected [3]. We used this information to find the optimal parameters to decompose an image into superpixels using SLIC [8]. Indeed, an error measure was estimated by counting how many superpixels were crossed by lines of multiple classes (Figure 5, B). Then, the compactness parameter of SLIC was optimized by minimizing the error estimate using sequential search.

**Figure 5.**
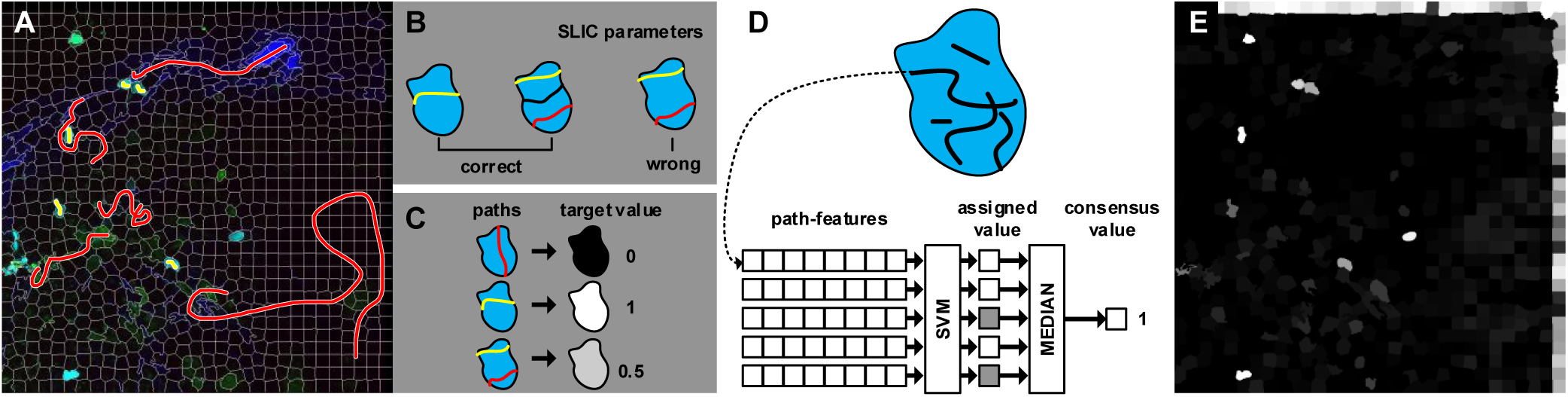
Path-based features for superpixel classification. **A.** 2P-IVM micrograph showing CFP neutrophils (light blue) in the popliteal lymph node of a CD11c/GFP animal (green) with collagen structures (blue - SHG) and viral particles (red - DiD). Yellow and red lines are the annotations provided by the users for disired and undesired points rpespectively. White lines are the borders of superpixel decomposition **B** The correctness of a superpixel decompiosition is evaluated by minimizing the number of lines of different class (yellow, red) in the same superpixel. **C.** Each superpixel is associated to a target value as the weighted average of the contained annotations. **D.** Path-based superpixel descriptor. (up) a superpixel is described as a set of paths. (low) each path is classified independently. then, the superpixel is classified as the mean of the classified paths. **E.** Output of superpixel classification, showing the disappearance of superpixels associated with cell debris (blebs). Artifacts on the border are related to a bug in the current version which will be fixed in the next preprints

A coarse decomposition on images into superpixels allowed to classify superpixels instead of single pixels, benefiting of the superpixel properties of boundary adherence and faster execution. The desired class (or target brightness value when using a linear prediction) can be estimated using the annotations provided by the user. If a superpixel is crossed by a line of desired objects, then its target value was set to 1. If a superpixel is crossed by a line of undesired objects, then its target value was set to 0. If a superpixel is crossed by multiple lines of different classes, then its value corresponds to the weighted average of the different classes (Figure 5, C).

Considering an image as a graph, having pixels as vertices and color differences as edges, shortest-paths on the image are useful to analyze its content [9, 3]. To this end, we described each superpixel as a set of shortest-path between arbitrary points inside the superpixel (Figure 5, D, up). A support vector machine was then trained to classify each path independently. The final predicted class for each superpixel was given by a consensus voting on the classes of the paths inside the superpixel, using, for instance, the median of the predicted value (Figure 5, D, low).

This method allowed to distinguish between superpixels containing cells and particles of disrupted cells (Figure 5, E).

Note for the preprint version: Figure 5, E reports the results from an older version of the software, with a bug that creates artifacts on the borders and less accuracy. These results are planned to be updated in the next revised version.

## Discussion

The proposed method relies on brushing, in line with other tools such as Ilastik [10] and Trainable WEKA Segmentation [11], asking the user to hand-draw some lines on interesting points (i.e. on some cells of interest) and other lines on non-interesting points (i.e. other cells, background, etc.). By contrast to the aforementioned tools, the proposed method exploits the provided information differently. More precisely, the annotations provided by the user (which are in the form of a line), are used to build a graph-based representation of the microscopy data.

Such a graph is used by the proposed method to compute specific features to face the challenges of 2P-IVM videos of immune cells.

Then, machine learning is applied on the graph to find a mathematical model that maps every single pixel to a scalar value, expressing how much it is likely to be part of the generated imaging channel. Once this model has been obtained, it can be applied to the entire video. Considering annotations from multiple videos, the method has the potential to be applied to process multiple videos without requiring parameters tuning.

Recently proposed approaches based on neuronal networks with convolutional layers such as U-NET [12] achieved remarkable performances for cell detection in microscopy data, using only limited annotations from the user. With respect to these approaches, the proposed method uses hand-crafted features. Although this choice is less generic and might not be optimal, it allows the user to choose the more appropriate features to face specific challenges in 2P-IVM imaging of the immune system.

## Methods

### Intravital two-photon microscopy

Deep tissue imaging was performed on a customized up-right two-photon platform (TrimScope, LaVision BioTec). Two-photon probe excitation and tissue second-harmonic generation (SHG) were obtained with a set of two tunable Ti:sapphire lasers (Chamaleon Ultra I, Chamaleon Ultra II, Coherent) and an optical parametric oscillator that emits in the range of 1,010 to 1,340 nm (Chamaleon Compact OPO, Coherent), with output wavelength in the range of 690–1,080 nm.

### Mice

All animals were bred in-house or acquired from Janvier labs (C57BL/6). Mice were maintained under specific pathogen-free conditions at the IRB, Bellinzona and used in accordance with the Swiss Federal Veterinary Office guidelines. CD11c-GFP [6], CK6/ECFP [5]. All animal experiments were performed in accordance with the Swiss Federal Veterinary Office guidelines and authorized by the relevant institutional committee (Commissione cantonale per gli esperimenti sugli animali, Ticino) of the Cantonal Veterinary with authorization numbers TI28/17, TI02/14 and TI07/13.

### Pixel-wise classification

Each pixel is classified as foreground or background, based on a predictive model trained on a few landmarks provided by the users. The predictive model is generated by training a Supported Vector Machine binary classifier using the radial basis function kernel. color features of each pixel, and in a circular neighborhood were used.

### Superpixel-wise classification

Videos were initially decomposed in superpixels using SLIC [8]. SLIC was applied on each slice of the 3d stack independently. A fixed number of 1000 superpixels was used for the results reported in this preprint version. Compactness was automatically determined by the method, minimizing the number of paths of different classes in each superpixel. Each superpixel was described employing K = 5 paths. Each path was described as the color-sequence of the vertices. Paths shorter, or longer than 8 points, were interpolated to 8 points using average smoothing.

### Implementation

The proposed method was entirely written in Matlab (Mathworks) as an extension for the bioimaging software Imaris and tested on the versions 7.7.2 to 9.3.1.

## Acknowledgments

We are thankful to Farsakoglu Y., I. Latino, T. Virgilio, M. Palomino-Segura for support in the generation of imaging data and for having tested the software. A. Pulfer, D. Morone, B. Thelen, G. Rovi, L. Karagyaur, A. Arini for technical support. This work was supported by the Swiss National Foundation (SNF) grants, 176124, R’equipt (145038) and Ambizione (148183), the European Commission Marie Curie Reintegration Grant (612742), the Center for Computational Medicine in Cardiology (CCMC) and SystemsX.ch for a grant to D.U.P. (2013/124).

The authors declare no competing financial interests.

## References

[1] Fritjof Helmchen and Winfried Denk. Deep tissue two-photon microscopy. Nature methods, 2(12):932–940, 2005.

[2] Jens V. Stein and Santiago F Gonzalez. Dynamic intravital imaging of cell-cell interactions in the lymph node. Journal of Allergy and Clinical Immunology, 139(1):12–20, 2017.

[3] Diego Ulisse Pizzagalli, Santiago Fernandez Gonzalez, and Rolf Krause. A trainable clustering algorithm based on shortest paths from density peaks. Science Advances, 5(10), 2019.

[4] Diego Ulisse Pizzagalli, Yagmur Farsakoglu, Miguel Palomino-Segura, Elisa Palladino, Jordi Sintes, Francesco Marangoni, Thorsten R. Mempel, Wan Hon Koh, Thomas T. Murooka, Flavian Thelen, Jens V. Stein, Giuseppe Pozzi, Marcus Thelen, Rolf Krause, and Santiago Fernandez Gonzalez. Leukocyte Tracking Database, a collection of immune cell tracks from intravital 2-photon microscopy videos. Scientific Data, 5:1–13, 2018.

[5] Anna-Katerina Hadjantonakis, Suzanne Macmaster, and Andras Nagy. Embryonic stem cells and mice expressing different GFP variants for multiple non-invasive reporter usage within a single animal. BMC biotechnology, 2(1):11, 2002.

[6] Peter B Stranges, Jessica Watson, Cristie J Cooper, Caroline-Morgane Choisy-Rossi, Austin C Stonebraker, Ryan A Beighton, Heather Hartig, John P Sundberg, Stein Servick, Gunnar Kaufmann, et al. Elimination of antigen-presenting cells and autoreactive t cells by fas contributes to prevention of autoimmunity. Immunity, 26(5):629–641, 2007.

[7] Ho Lee, Clemens Alt, Costas M Pitsillides, Mehron Puoris’haag, and Charles P Lin. In vivo imaging flow cytometer. Optics express, 14(17):7789–7800, 2006.

[8] Radhakrishna Achanta, Appu Shaji, Kevin Smith, Aurelien Lucchi, P Fua, and S Susstrunk. SLIC Superpixels. EPFL Technical Report 149300, (June):15, 2010.

[9] Stefano Ghidoni, Loris Nanni, Sheryl Brahnam, and Emanuele Menegatti. Texture descriptors based on dijkstra’s algorithm for medical image analysis. In Studies in health technology and informatics, pages 74–82. IOS, 2014.

[10] Christoph Sommer, Christoph Straehle, Ullrich Köthe, and Fred A Hamprecht. Ilastik: Interactive learning and segmentation toolkit. In 2011 IEEE international symposium on biomedical imaging: From nano to macro, pages 230–233. IEEE, 2011.

[11] Ignacio Arganda-Carreras, Verena Kaynig, Curtis Rueden, Kevin W Eliceiri, Johannes Schindelin, Albert Cardona, and H Sebastian Seung. Trainable Weka Segmentation: a machine learning tool for microscopy pixel classification. Bioinformatics, 33(15):2424–2426, 2017.

[12] Thorsten Falk, Dominic Mai, Robert Bensch, Özgün Çiçek, Ahmed Abdulkadir, Yassine Marrakchi, Anton Böhm, Jan Deubner, Zoe Jäckel, Katharina Seiwald, et al. U-net: deep learning for cell counting, detection, and morphometry. Nature methods, 16(1):67, 2019.

